# Loss of malate-aspartate shuttle component SLC25A12 induces pulmonary metastasis

**DOI:** 10.1101/2020.05.29.121400

**Authors:** H Furkan Alkan, Paul W Vesely, Hubert Hackl, Johannes Foßelteder, Matthew G Vander Heiden, Martin Pichler, Gerald Hoefler, Juliane G Bogner-Strauss

## Abstract

**Background:** Aspartate biosynthesis and its delivery to the cytosol can be crucial for tumor growth *in vivo.* However, the impact of aspartate synthesis on metastasis has not been studied. We previously described that loss-of-aspartate glutamate carrier 1 (SLC25A12 or AGC1), an important component of the malate-aspartate shuttle, impairs cytosolic aspartate levels, NAD^+^/NADH ratio, mitochondrial respiration, and tumor growth. Here, we report the impact of AGC1-knockdown on metastasis.

**Results:** AGC1 expression is positively correlated with worse patient prognosis in many cancers. AGC1-knockdown in mouse lung carcinoma and melanoma cell lines leads to increased pulmonary metastasis following subcutaneous or intravenous injections, respectively. On the other hand, conventional *in vitro* metastasis assays show no indication of increased metastasis capacity of AGC1-knockdown cells.

**Conclusion:** This study highlights that certain branches of metabolism impact tumor growth and tumor metastasis differently. In addition, it also argues that commonly known metastasis indicators, including EMT genes, cell migration, or colony formation do not always reflect the metastatic capacity *in vivo*.

## Introduction

Tumor metastasis is correlated with poor prognosis. Surgical removal of the metastatic tumors from one or more organs is challenging and drug-resistance-induced by the new tissue environment-may occur in metastatic tumors (Fendt, 2019). To form metastasis, cancer cells follow a cascade of events including invading the surrounding tissue, detaching from the primary tumor, surviving in the circulation, and colonizing a distant organ (Elia et al., 2018; Peiries-Peres et al., 2015). In addition to the bioenergetics and biosynthetic demands of cell proliferation, metastatic cells also need to adapt their metabolism to succeed in the different steps of this cascade of events (Fendt, 2019). Although a definitive metabolic pattern that distinguishes metastatic from non-metastatic tumors has not yet been identified, there is evidence that certain metabolic pathways could be more important for metastasis formation (Elia et al., 2018). For instance, antioxidants such as N-acetylcysteine (NAC) increase melanoma metastasis in mice (Le Gal et al., 2015; Peiries-Peres et al., 2015), suggesting that suppressing oxidative stress is one requirement of distant metastasis. Consistently, inhibition of the folate pathway using methotrexate or knockdown of either ALDH1L2 (Aldehyde Dehydrogenase 1 Family Member L2) or MTHFD1 (Methylenetetrahydrofolate Dehydrogenase) blocks melanoma metastasis *in vivo* via hindering the survival of circulating melanoma tumors while having no impact on subcutaneous tumor growth (Piskounova et al., 2015). In addition, proline catabolism is selectively important for the proliferation and colony formation of metastatic cells and reduced Prodh (proline dehydrogenase) activity leads to impaired *in vivo* lung metastasis in mouse breast cancer tumors (Elia et al., 2017). Pyruvate was also shown to be an important mediator of metastatic niche formation through hydroxylation of collagen (Elia et al., 2019). Furthermore, asparagine levels or ASNS (asparagine synthetase) activity could also determine the metastatic capacity of breast cancers both *in vivo* and *in vitro* (Knott et al., 2018).

Aspartate is a proteinogenic non-essential amino acid that is used for de novo synthesis of nucleotides and certain non-essential amino acids. Aspartate is also involved in several intracellular processes including, but not limited to, redox homeostasis and the urea cycle. Intracellular aspartate levels correlate with the proliferation rate in cancer cells (Gui et al., 2016) and are limiting for the growth of some tumors *in vivo* (Garcia-Bermudez et al., 2018; Sullivan et al., 2018). Although aspartate is available in circulation, these levels are insufficient to trigger its uptake in most cells that lacking the plasma membrane aspartate transporter SLC1A3 (Garcia-Bermudez et al., 2018). Therefore, endogenous aspartate synthesis can limit the growth of some tumors. For example, one requirement for aspartate production is a suitable NAD^+^/NADH ratio. Cells lacking mitochondrial respiration display plummeting aspartate levels and slow proliferation (Sullivan et al., 2015; Birsoy et al., 2015). In addition, glutamine is an important substrate for mitochondrial aspartate synthesis. In our previous study, we determined that blocking glutamine metabolism and/or inhibiting cytosolic aspartate delivery via knockdown of mitochondrial aspartate-glutamate carrier 1 (AGC1) impairs cell proliferation and tumor growth (Alkan et al., 2018). Altogether, our previous findings suggest that not only having the adequate conditions (high NAD^+^/NADH ratio) and the substrate (glutamine) available for aspartate synthesis but also the ability of the mitochondria to export its aspartate to the cytosol can be a limitation for cell proliferation (Alkan and Bogner-Strauss, 2019).

In this follow-up study, we report that AGC1 knockdown (KD) in mouse lewis lung carcinoma (LLC1) cells leads to increased lung metastasis and poor overall survival of syngeneic mice, despite impaired subcutaneous tumor growth. Similarly, we observed enhanced metastatic spread of intravenously injected AGC1-KD B16F10 cells upon AGC1-KD. In addition, conventional *in vitro* metastasis assays show no indication of increased metastasis capacity of AGC-KD cells, suggesting that the role of aspartate metabolism in metastasis formation is relevant to metastatic events occurring only *in vivo*. Collectively, our report argues that certain branches of metabolism impact tumor growth and tumor metastasis differently.

## Materials & Methods

### Bioinformatics

Gene expression profiles (RNA sequencing V2 data) of SLC25A12 (Aralar or AGC1) and SLC25A13 (Citrin or AGC2), as well as clinical information for 21 different cancer type cohorts of The Cancer Genome Atlas (TCGA) were retrieved via firebrowse.org (courtesy Broad Institute of MIT & Harvard). Survival analyses were performed on dichotomized groups of patients with high and low expression levels using an optimized cut-off according to the maximal concordance index. The Kaplan Meier method was used to estimate survival distributions and the log-rank test was used to evaluate statistical significance in OS between patient groups. Where stated, p-values were adjusted for multiple testing as described previously (Altman et al., 1994). Log2 hazard ratios and 95% confidence interval were visualized as forest plot. Differences in gene expression of the primary tumor between pathological M stages were tested using the none-parametric two-sided Wilcoxon rank sum test and visualized as boxplots. All analyses were performed using R and packages survival and dynpred.

### Cell Culture

All cells used were tested negative for Mycoplasma. All cell lines were maintained in DMEM (25 mM Glucose, 4 mM Glutamine, 1 mM Sodium Pyruvate) supplemented with 10% Fetal Bovine Serum (FBS, Gibco) and 50 units/mL Penicillin/Streptomycin (Gibco) in sterile incubators set to 37°C temperature and 5% CO_2_ concentration. AGC1 protein was knocked-down using shRNA as described previously (Alkan et al., 2018).

### Scratch Assay

6×10^5^ cells were seeded in a 6-well plate and incubated for 24 hours. Cells were harvested by trypsinating and 1×10^5^ transfected cells were seeded in a 24-well culture plate in 6 replicates and incubated for 24 hours to form a confluent cell monolayer. The scratch was introduced using a 200μL pipette tip and detached cells were washed off with PBS.

Cells were further incubated with fresh growth media and scratch closure was observed using an inverted light microscope (Olympus IX71). The scratch width was measured using the cellSens imaging software (Olympus) every 4-12h depending on the cell line. Arithmetic mean of all replicates was used to calculate the scratch closure of cells.

### Sphere Formation Assays

To test the ability of cells with modified gene expression levels to form tumor spheres, we generated tumor spheres as previously described (Schwarzenbacher et al., 2019). In detail, the adherent growing cell lines were dissociated into single cells using trypsin/EDTA and 2.000 single cells per well seeded in ultra-low attachment 6-well plates (Corning, NY, USA) using serum-free MEBM (Lonza, Basel, Switzerland) medium (SFM) supplemented with 1xB27 supplement (Gibco), 20 ng/ml human epidermal growth factor EGF (Peprotech, Hamburg, Germany), 10 ng/ml human basic fibroblast growth factor FGF (Peprotech), 20 IU/ml Heparin (Baxter, Vienna, Austria) and 1 % antibiotic/antimycotic solution (Thermo Fisher Scientific, containing 10,000 units/mL of penicillin, 10.000 μg/mL of streptomycin, and 25 μg/mL of Gibco Amphotericin B). After 14 days, the number of spheres was counted using a bright-field microscope in three independent replicates.

### Migration Assay

5000 cells in 100μL DMEM containing 1% FBS were seeded into insert wells of the Transwell plates (Corning). Lower wells were filled with 500μL DMEM containing 10% FBS. 48 hours later, cell numbers in both upper and lower wells were measured via either conventional cell counting or using Cell Titer Glo (Promega) as described previously (Alkan et al., 2018).

### ROS Measurement

Sub-confluent LLC1 cells were incubated with 1 μM CellRox (RED2 Channel, Invitrogen) solution for 1 hour, washed twice with PBS, trypsinized, and resuspended with media by pipetting long enough to obtain single cells. After centrifugation, cells were mixed well with 100 μL PBS and incubated with 1:0000 Ghost Dye (NIR2 Channel, Tonbobio) for half an hour. Ghost Dye was washed away and cells were resuspended in PBS. Fluorescent signals were obtained via Guave Easy Cyte 8 (Millipore) and data were analyzed using InCyte Software (Millipore).

### Animal Care

The animal study was approved by the institutional ethics committee and experiments were performed according to the guidelines of the Austrian Federal Ministry of Science and Research. Experiment licenses were granted under BMWF-66.007/0026-WF/V/3b/2015 and BMWF-66.007/0008-WF/V/3b/2016. In vivo, CB-839 experiments were approved by the MIT Committee on Animal Care (IACUC).

LLC1 cells were cultured in T175 flasks about a week prior to the day of injection. For getting ready with the injections, cells were grown to 50-60% confluency, washed with PBS, mixed with 5 mL trypsin and resuspended with 25 mL media as soon as they detach from the plate. Cells were flown through 0.45 mm × 25 mm needles to avoid clumps; and centrifuged for 5 min at 1000 rpm. Media was aspirated, the pellet was resuspended with 3 mL ice-cold HBSS, viable cells were counted using a hemocytometer and diluted to 2.5×10^6^ cells/mL in HBSS. Cells were kept at 4°C and used for injection within less than 2 hours. 0.5×10^6^ cells (200μL) were injected over the right flanks or behind the neck of 8-week-old female C57BL/6 mice (8 mice per group) using 0.6 mm × 30 mm needles. Mice were sacrificed at day 16 or day 21 and tumor and lung tissues were collected for further analysis. The survival rate of the mice was monitored at least once every day in standard housing conditions ensuring their access to water and food supply.

B16F10 cells were cultured and prepared for injection as it is described above for LLC1 cells. Cells were diluted to 3×10^6^ cells/mL in cold HBSS. 8-10 week-old female C57BL/6 mice were restrained leaving only their tail outside, providing access to open-air through small holes near face area and 0.3×10^6^ cells (100μL) were injected into the tail vein using 0.3 mm × 12 mm needles (Omnican). The mice were harvested at day 16 and lungs were collected, rinsed, and fixed in 10% formalin for at least 48 hours.

For pulmonary tumor area measurement, each lung was sliced into 12-15 pieces, embedded in paraffin, stained with Hematoxylin and Eosin (H&E), and total area, as well as tumor area, was measured for every piece using NisElements Software. Tumors were determined at 10X magnification; areas were measured at 2X.

### Quantitative Real-Time PCR

Total RNA from cells was isolated using the PeqGOLD total RNA isolation kit (Peqlab). Tissue RNA was isolated with TRIzol reagent (Life Technologies). cDNA was generated using the Qiagen QuantiTect RT kit. mRNA expression was assessed using SYBR green real-time PCR on an ABI 7000 instrument as described. Unless stated otherwise, Ct values of every gene first subtracted from the ones of Rplp0 (for LLC1 cells) and TfIIb (for B16F10 cells) within every cDNA sample, then normalized to the mean δCt of the control group for every gene. Fold changes were calculated by taking the δδCt power of (½).

### Statistical analysis

All experiments were performed using at least three independent replicates (different passages and mice for *in vitro* and *in vivo* experiments, respectively). Single comparison between two groups, is calculated via two-tailed, non-parametric, unpaired student’s t-test. Significance levels: * p ≤ 0.05, ** p ≤ 0.01, *** p ≤ 0.001.

## Results

### Low AGC1 expression correlates with worse prognosis in some cancers

We previously generated shRNA-mediated knockdowns (KD) of mitochondrial aspartate-glutamate carrier 1 (AGC1) and observed that AGC1-KD inhibited Lewis Lung Carcinoma (LLC1) tumor growth (Alkan et al., 2018). However, mice with AGC1-KD tumors tended to die earlier than controls, even though their tumors were smaller (Figure 1A,1B). This implies that AGC1 might have an impact on cancer beyond affecting cell proliferation and tumor growth. Analysis of overall patient survival and gene expression from The Cancer Genome Atlas (TCGA) showed that human lung adenocarcinoma tumors (LUAD) with lower AGC1 (SLC25A12) mRNA expression also correlated with worse prognosis (Figure 1C). Although this correlation was not universal across all cancer types analyzed, several other cancer including Cervical Squamous Cell Carcinoma (CESC), Kidney Renal Clear Cell Carcinoma (KIRC), Kidney Renal Papillary Cell Carcinoma (KIRP), and Sarcoma (SARC) showed a similar association between AGC1 expression and worse overall patient outcome (Figure 1D). Interestingly, the correlation between expression of the functionally identical AGC2 (SLC25A13) isoform (Thangaratnarajah et al., 2014) and patient survival was usually in the opposite direction of AGC1 within the same type of cancer (Figure 1D). Only in kidney renal cell carcinomas, low expression levels of both AGC1 and AGC2 showed a distinct correlation with the poor prognosis of patients (Figure 1D,E). Further bioinformatic analysis of this type of cancer suggested mRNA expressions of both AGC1 and AGC2 were lower in metastatic regions compared to the primary tumors (Figure 1F), implying that AGC1 might influence metastatic spread in human cancers.

**Figure 1:**
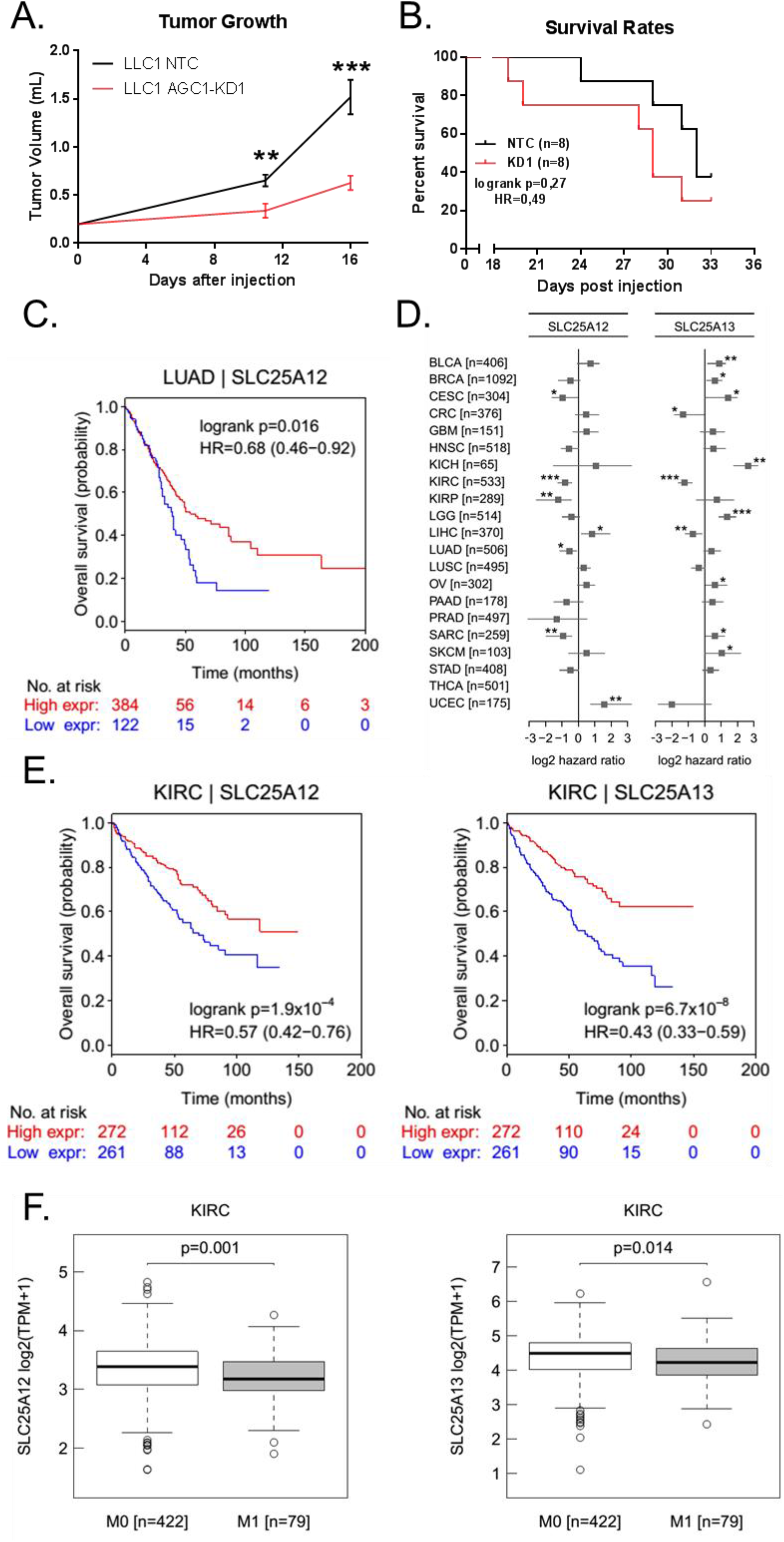
AGC1-deficient tumors tend to be more lethal for mice. (A) Tumor progression of control (NTC) and AGC1-KD LLC1 tumors measured as described above. (B) Survival rates of the mice bearing control (NTC) and AGC1-KD LLC1 tumors monitored over the course of 33 days. (C) Overall survival rates of patients with high (red line) or low (blue line) AGC1 (SLC25A12) mRNA-expressing lung adenocarcinomas (LUAD). Data is adopted from the TCGA database. Patients with higher or lower AGC1 (SLC25A12) mRNA expression was divided into two groups with a cut-off line where the separation of these groups was significantly most meaningful. (D) Correlation of AGC1 (SLC25A12) or AGC2 (SLC25A13) mRNA expression with better (negative log2 hazard ratio) or poor (positive log2 hazard ratio) survival of patients from various cancers. Lower log2 hazard ratio for a gene means that the group of patients with low expressing tumors have worse overall survival than the high expressing group. The cut-off to separate groups was determined as described in C. (E) Overall survival (OS) rates of patients with high (red line) or low (blue line) AGC1 (SLC25A12) or AGC2 (SLC25A13) mRNA-expressing kidney renal clear cell carcinoma tumors (KIRC). Data is adopted from the TCGA database. The cut-off to separate groups was determined as described in C. (F) mRNA expressions of AGC1 (SLC25A12) or AGC2 (SLC25A13) in kidney renal clear cell carcinoma tumors (KIRC) of patients without (M0) or with metastasis (M1).

### AGC1-KD increased the pulmonary metastasis from LLC1 and B16F10 cells

Based on the bioinformatics analyses and because metastasis is the leading cause of death in cancer patients, we hypothesized that AGC1-deficient cells might have a superior metastatic capacity compared to control cells. Thus, we performed a second line of subcutaneous LLC1 tumor cell injections and assessed the lungs of the mice at the end of the experiment. Strikingly, histological analysis showed that none of the mice from the control group had pulmonary metastasis while approximately half of the mice bearing AGC1-KD tumors had evidence of lung metastasis (Figure 2A). To achieve bigger primary tumors and more metastasis, we next injected LLC1 cells subcutaneously behind the neck of the mice instead of the flanks. This approach allows injecting higher number of cells and a long period of tumor growth. Eventually, we sacrificed the mice and harvested the lungs 21 days after tumor cell injection, and the percentage of tumor area within the lung was measured by morphometric analysis. We observed the metastatic regions within the lungs from the mice bearing AGC1-KD LLC1 tumors to be significantly larger than those from mice bearing control tumors (Figure 2B). Because AGC1-depletion synergizes with glutaminase inhibition (Alkan et al., 2018), we tested whether mice treated with the glutaminase inhibitor CB-839 would have a different metastasis outcome. However, inhibiting glutamine metabolism *in vivo* had no impact on metastasis of either control or AGC1-KD tumors (Figure 2C).

**Figure 2:**
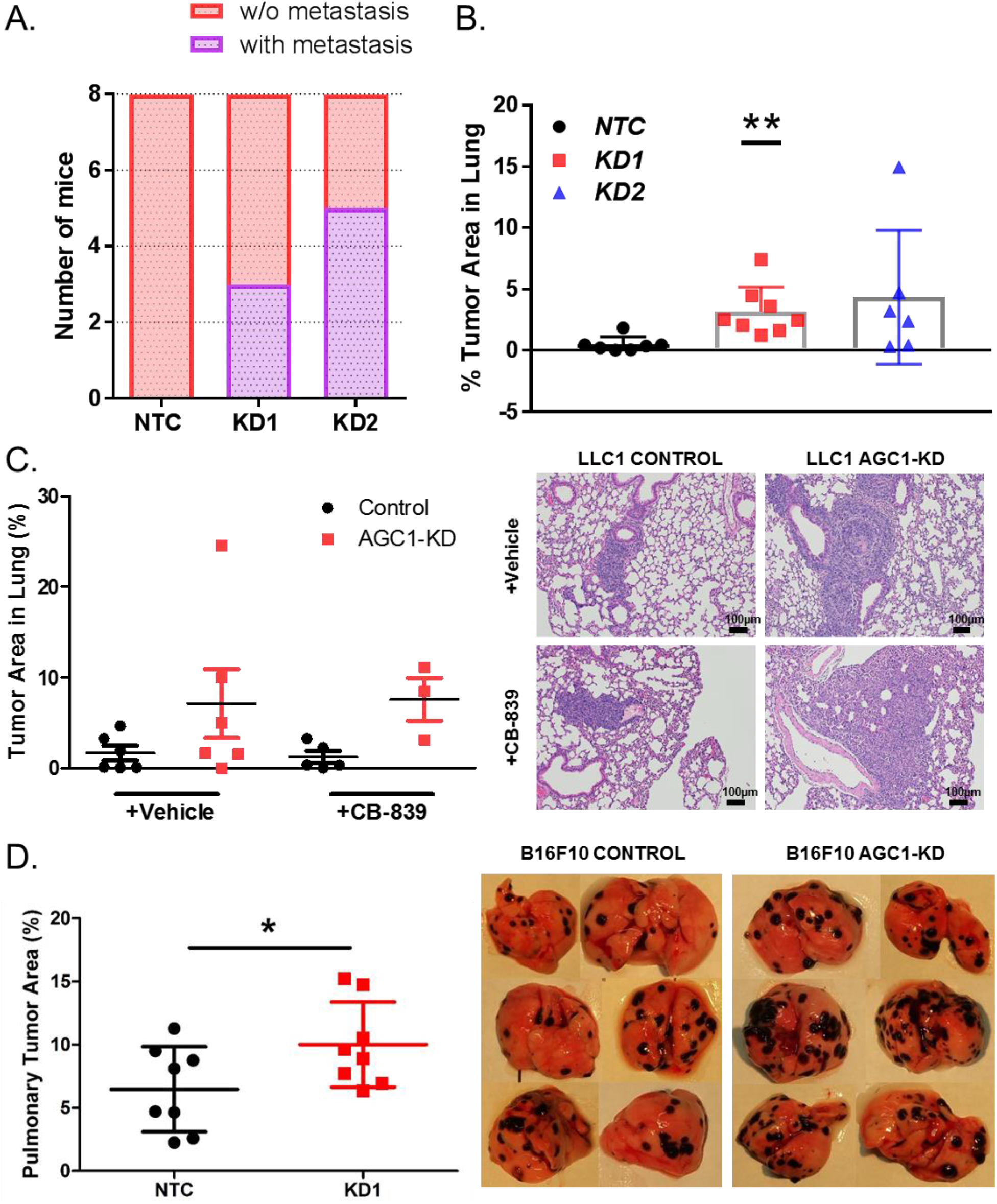
Cancer cells with AGC1-knockdown have higher potential to metastasize. (A) The number of mice with (purple) or without (pink) lung metastasis 16 days after control (NTC), or AGC1-KD LLC1 cells (KD1, sh911 or KD2, sh908) were injected in the flanks. H&E stained lung slides were screed for metastasis. (B) Percent of the metastatic area in the lungs of mice bearing control (NTC), or AGC1-KD (KD1 with sh911; KD2 with sh908) LLC1 tumors, measured 21 days after cells were injected subcutaneously behind the necks of mice (n≥6). Lungs were resected and H&E stained sections were analysed (C) (left) Percent of the metastatic area in the lungs of mice that were bearing control (NTC), or AGC1-KD LLC1 tumors on the flanks 22 days after injections. Mice were treated without (Vehicle) or with CB-839 dosed at 200 mg/kg twice daily starting on day 13 (n≥6). (right) Representative histology pictures of the metastatic areas of mice injected with control (NTC) or AGC1-KD LLC1 tumors. (D) (left) Percent of the metastatic areas (%) in the lungs of mice with control (NTC), or AGC1-KD LLC1 cells, measured 16 days after intravenous injection of cells (n=8). (right) Representative images of lungs with metastasis harvested 16 days after control (NTC), and AGC1-KD B16F10 cells were intravenously injected. Significance levels: * p ≤ 0.05, ** p ≤ 0.01, *** p ≤ 0.001

To test whether AGC1-loss also increases metastasis formation in a different cancer model, we knocked-down AGC1 in B16F10 mouse melanoma cell line that is specifically generated for their metastatic potential (Hart, 1979). AGC1 depletion in B16F10 cells reduced cell proliferation to a comparable extent as in LLC1 cells and in other cell lines from our previous study (Figure S1A) (Alkan et al., 2018). To test the impact of AGC1-knockdown on the metastatic spread of B16F10 cells *in vivo*, we injected them intravenously in mice. 16 days post-injection, mice were sacrificed, lungs were harvested and the percentage of tumor area within the lung was determined as described above. Strikingly, AGC1-KD B16F10 cells also yielded more metastatic areas in the lungs compared to control cells (Figure 2D, S1B). These findings suggest that AGC1-loss is correlated with increased metastatic capacity in independent cancer models.

### Increased metastatic capacity of AGC1-KD cells cannot be detected by *in vitro* methods

The finding that the AGC1-KD tumor cells show relatively slow proliferation but high metastatic potential led to the conclusion that biological events other than proliferation rate likely to cause this phenomenon. Hence, we performed a series of *in vitro* tests to analyze migration speed and colony formation capacity. First, we performed a scratch (wound-healing) assay to assess the ability of AGC1-KD cells to migrate. However, we found that AGC1-KD LLC1 and B16F10 cells covered the scratched area slower compared to corresponding control cells (Figure 3A,3B). Because the outcomes from this experiment may be due to the reduced proliferation rate of AGC1-KD cells, we next performed migration assays using trans-well plates. We normalized the number of cells that migrated into the bottom plate to the proliferation rate and still found that the migration rate of AGC1-KD LLC1 cells was almost identical to those of control cells (Figure 3C). Next, we measured three-dimensional tumor sphere formation ability of AGC1-KD cells. Consistent with proliferation, migration, and wound-healing assays, AGC1-KD cells showed a reduced capacity to form spheres under low-attachment conditions (Figure 3D,3E). Only in B16F10 cells, the sizes of the tumor spheres of AGC1-KD cells were comparable to the colony sizes of the control cells, although this also did not explain the increased metastatic phenotype observed *in vivo* (Figure 3F,3G). In summary, AGC1-KD cells showed no evidence of improved metastatic capacity using common surrogate assays for metastatic cell properties *in vitro*.

**Figure 3:**
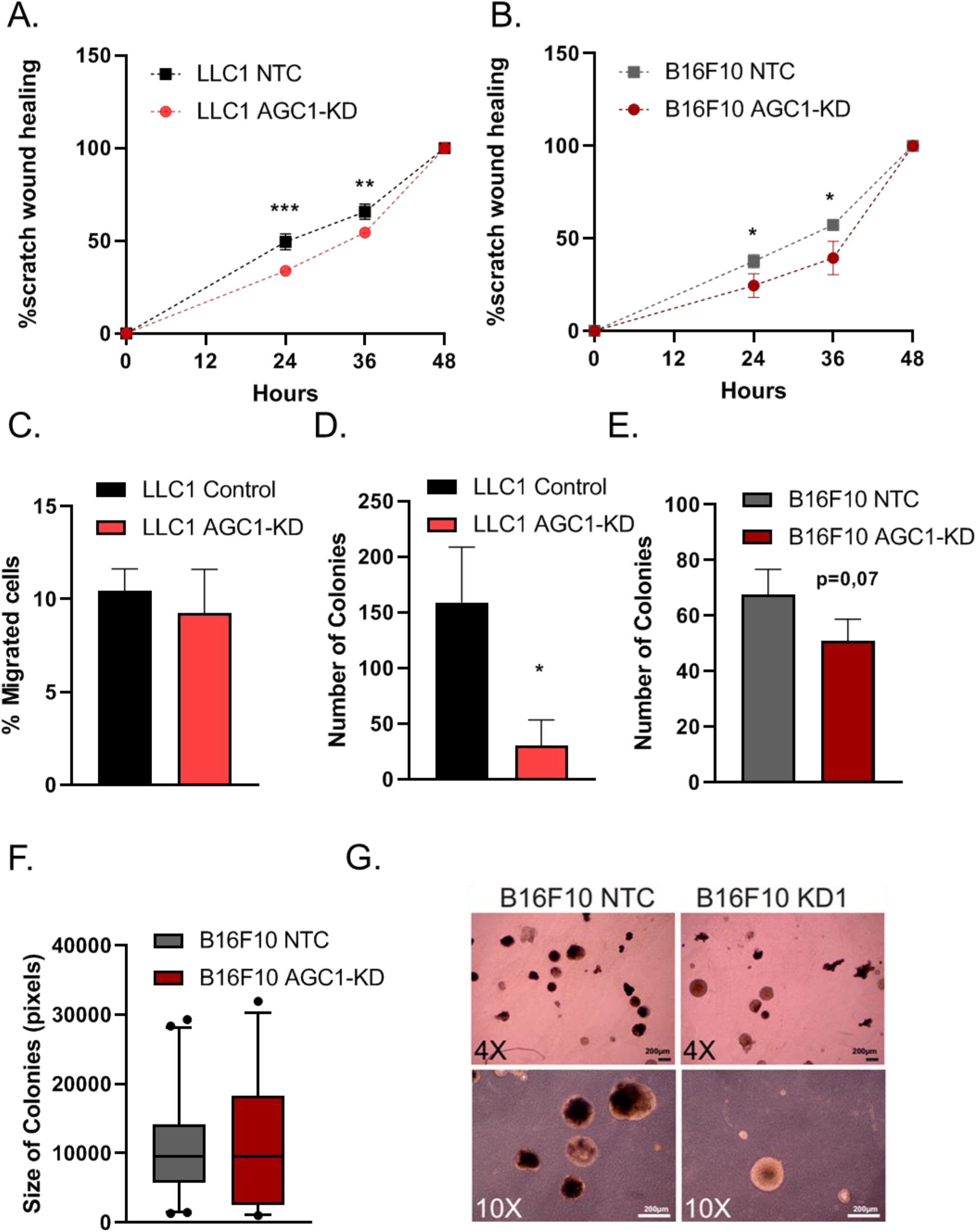
AGC1-KD cells do not show signs of increased metastatic ability in cell culture. (A-B) Percent area covered by control (NTC) and AGC1-KD (A) LLC1 or (B) B16F10 cells over the course of 24, 36, and 48 hours following the scratching (n=5). (C) Percent of control (NTC) and AGC1-KD cells migrated to the bottom wells of the trans-well after 48 hours (n=3). (D-E) The number of spheres 14 days after seeding of 2.000 control (NTC) and AGC1-KD LLC1 and B16F10 cells, counted by bright field microscopy (n=3). (F) Average colony sizes of B16F10 cells from the sphere formation assay in E. (G) Representative picture of spheres from B16F10 cells. All experiments denote mean ±SDs. Significance levels: * p ≤ 0.05, ** p ≤ 0.01, *** p ≤ 0.001

### Genes related to Aldh1l1 metabolism are deregulated in AGC1-KD cells

Epithelial-to-mesenchymal-transition (EMT) is an important hallmark of cancer metastasis. Accordingly, it is thought that it is a pre-requisite for cancer cells to switch from an epithelial-to a mesenchymal-cell surface phenotype to allow for escape from the primary tumor tissue mass (Elia et al., 2018). To test whether LLC1 tumors with AGC1-KD show an enhanced mesenchymal phenotype compared to controls, we measured mRNA expressions of common EMT and stemness marker genes in AGC1-KD tumors. Snai1, Snai2, Sox2, Skp2, and Twist1 were previously reported to induce EMT and metastasis, and E-cadherin expression is reversely correlated with EMT (Gao et al., 2015; Yang et al., 2014). None of these markers, however, showed expression differences between KD and control groups (Figure 4A). According to this definition, LLC1 AGC1-KD tumors did not show an increased EMT phenotype compared to controls.

**Figure 4:**
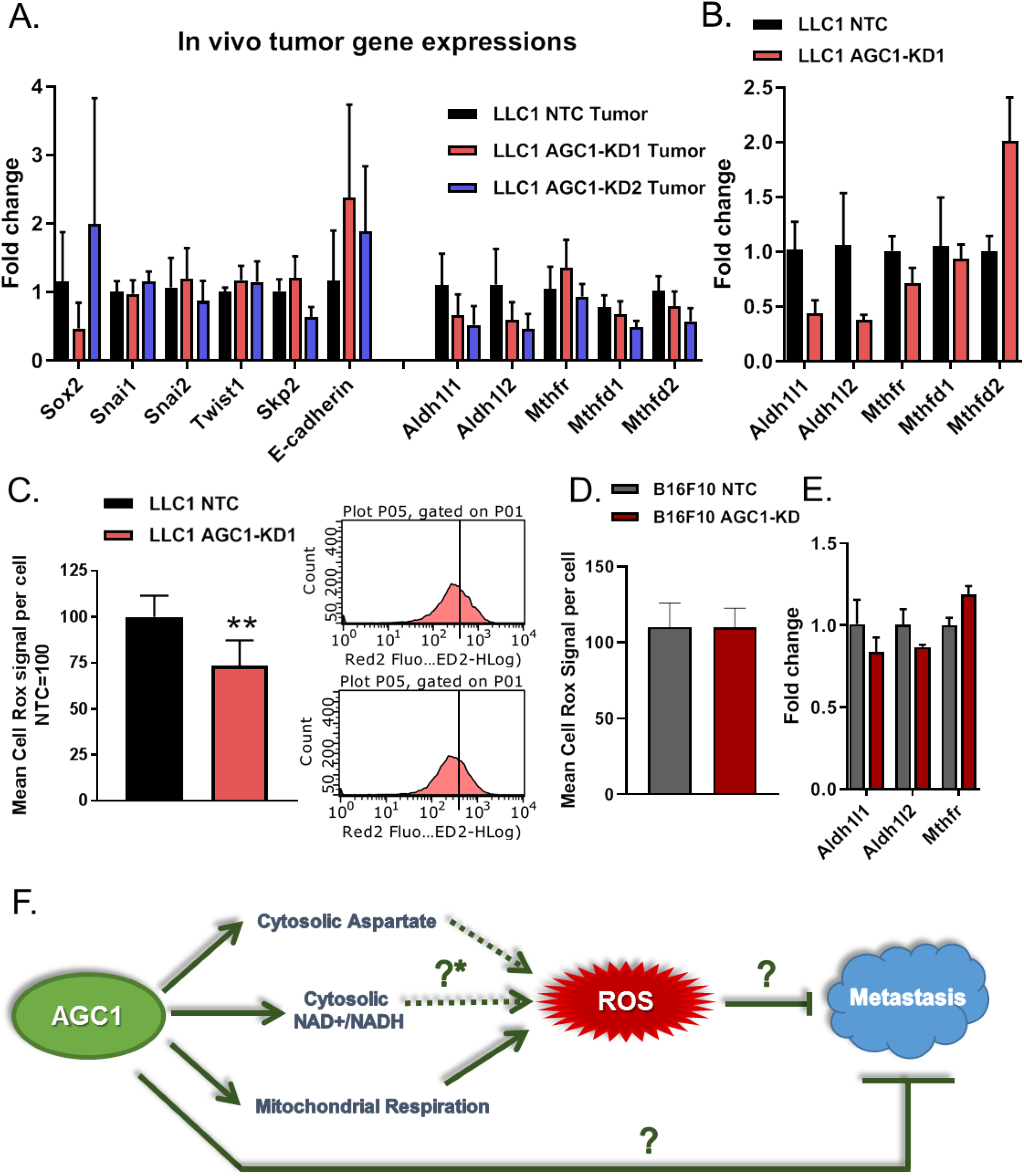
AGC1-KD LLC1 cells deregulate folate metabolism (genes) and have lower levels of ROS. (A) mRNA levels of EMT markers and folate pathway genes in control (NTC), AGC1-KD (KD1, sh911 and KD2, sh908) LLC1 tumors in *vivo* (n≥6). (B) mRNA levels of folate pathway genes in control (NTC), AGC1-KD1 LLC1 cells *in vitro* (n=3). (C) (left) Mean CellRox signal per well in AGC1-KD and control (NTC) LLC1 cells; normalized to controls (n=3). (right) histograms of one representative well from both control (NTC) and AGC1-KD LLC1 cells showing CellRox signal measured via flow cytometry (D) Mean CellRox signal per well in AGC1-KD and control (NTC) B16F10 cells; normalized to controls (n=3). (E) mRNA levels of folate pathway genes in control (NTC), AGC1-KD1 B16F10 cells *in vitro* (n=3). (F) Schematic demonstration of potential connections among AGC1 expression, intracellular ROS levels and metastasis. [?*] is referring to the potential changes in folate pathway upon AGC1 expression. All experiments denote mean ±SDs. Significance levels: * p ≤ 0.05, ** p ≤ 0.01, *** p ≤ 0.001

As a mediator of cellular oxidative stress, folate metabolism can be another important driver that dictates metastatic capacity (Singh et al., 2013; Piskounova, et al., 2015). Therefore, we measured gene expression levels of folate pathway enzymes in LLC1 tumor and cultured cells. Aldh1l1 and Aldh1l2 mRNA levels were consistently lower in AGC1-KD tumors and cultured cells compared to corresponding controls (Figure 4A, 4B), suggesting that folate pathway might be deregulated upon AGC1-loss. In addition, intracellular reactive oxygen species (ROS) levels were significantly lower in AGC1-KD LLC1 cells (Figure 4C). This is consistent with potentially altered folate metabolism and is consistent with the findings from our previous study that AGC1-KD reduced mitochondrial oxygen consumption (Alkan et al., 2018). Because suppressing oxidative stress can assist the survival of circulating cells (Piskounova, et al., 2015), lower baseline levels of ROS might be one explanation why AGC1-depletion has a positive impact on metastatic capacity. However, B16F10 cells showed no difference in ROS levels (Figure 4D) or folate-pathway genes upon AGC1-KD (Figure 4E) despite enhanced metastatic spread capacity (Figure 3D), arguing other pathways must also be involved (Figure 4F).

An ability to escape the immune response is important for cancer cells to survive in circulation and form metastasis in distant organs (Janssen et al., 2017; Blomberg et al., 2018). In our previous study, we found that AGC1-KD cells have lower cytosolic NAD^+^/NADH ratio and are less glycolytic (Alkan et al., 2018). Because inflammatory cells are often more glycolytic than anti-inflammatory cells (Neagu et al., 2019), we speculated that AGC1-KD cells might have an advantage to evade immune response by secreting anti-inflammatory cytokines. However, mRNA levels of tested cytokines were not significantly altered in AGC1-KD LLC1 tumors compared to the controls, although some anti-inflammatory cytokines such as Interleukin-10, Granulocyte-macrophage colony-stimulating factor (GM-CSF), and Transforming growth factor beta (TGFβ) showed tendencies towards upregulation in AGC1-KD tumors (Figure S2A). On the other hand, findings from the *in vitro* experiment and B16F10 cells were again inconsistent with this observation in LLC1 cells (Figure S2B,S2C). These findings argue that contributions of many pathways, or an unknown mechanism, is responsible for the metastatic capacity of AGC1-KD tumors *in vivo*.

## Discussion

In this study, we report that reduced mitochondrial aspartate export and malate-aspartate shuttle activity boost pulmonary metastasis of mouse lung (LLC1) and melanoma (B16F10) cell lines *in vivo*, despite slowing down cell proliferation and primary tumor growth (Alkan et al., 2018). To the best of our knowledge, this is the first report indicating that impaired aspartate metabolism, which is a limiting metabolite for tumor growth, increases the metastatic potential of cancer. Furthermore, this report highlights that commonly known metastasis indicators, including EMT genes, cell migration, or colony formation do not always reflect the metastatic capacity *in vivo* and our study provides strong evidence that metabolic requirements of metastasis may be different from proliferation itself.

It is important to note that the exact mechanism of how the loss of AGC1 induces pulmonary metastasis is not fully explained in this study. Altered levels of folate pathway genes, reduced intracellular ROS levels, the lack of enhanced colonization or migration phenotypes *in vitro* may indicate that AGC1-knockdown cells have an advantage surviving in the circulation (Piskounova, et al., 2015). Although AGC1-KD B16F10 cells do not have reduced baseline ROS levels *in vitro*, this might be different *in vivo*. Several other metabolites including pyruvate, proline, and asparagine were also reported to be involved in the metastasis process (Elia et al., 2019, Knott et al., 2019, Elia et al., 2017). However, we failed to detect any increase in these metabolites *in vivo* in AGC1-KD tumors during our previous study that could reflect such metastatic phenotype (Alkan et al., 2018). Of note, those studies mainly focused on breast cancer metastasis and lung cancer and melanoma cells may require different metabolic pathways to form metastasis.

AGC1 is an important mediator of the malate-aspartate shuttle, cytosolic aspartate levels, cytosolic NAD^+^/NADH ratio, and mitochondrial NADH levels (Alkan et al., 2018). Higher MCT1 expression and lactate consumption are correlated with efficient metastasis (Tasdogan et al., 2020). Reduced NAD^+^/NADH and pyruvate/lactate ratios in AGC1-KD cells (Alkan et al., 2018) may contribute to increased lactate uptake *in vivo*, leading to a similar metastatic phenotype reported in high MCT1-expressing cells. In addition, reduced mitochondrial respiration upon AGC1-knockdown can explain lower ROS production. Nevertheless, how folate pathway could be linked to aspartate transport or NAD^+^/NADH ratio needs to be studied further. Because the metastatic phenotype is not observed *in vitro*, we were limited in this study to further examine the mechanistic explanations of why AGC1-KD induces metastasis. However, these findings could lay the groundwork for future studies investigating metabolic requirements of cancer metastasis and how mitochondrial deficiencies may have adverse effects on tumor growth and metastasis.

## Supplemental Information

Supplemental Information includes 2 Supplemental Figures and their legends.

## List of Abbreviations

AGC1/2: Mitochondrial Aspartate-Glutamate Carriers ½
EMT: Epithelial-to-mesenchymal-transition
GM-CSF: Granulocyte-macrophage colony-stimulating factor
KD: Knockdown
NAC: N-acetylcysteine
NAD^+^/NADH: Nicotinamide adenine dinucleotide
ROS: Reactive Oxygen Species
TCA: Tricarboxylicacid cycle
TGFβ: Transforming growth factor beta

## Competing interests

The authors declare no competing interests.

## Available data and materials

The authors declare that all data supporting the findings of this study are available within the article and its supplementary information files and from the first and corresponding author upon reasonable request. Contact

## Funding

This work was funded by the Austrian Science Fund FWF SFB LIPTOX F3018, P27108, and W1226 DK “Metabolic and Cardiovascular Disease”. M.G.V.H. acknowledges support from the Lustgarten Foundation, SU2C, the Ludwig Center at MIT, the MIT Center for Precision Cancer Medicine, and the NCI (P30 CA1405141, R01 CA168653). M.G.V.H. is also an HHMI faculty scholar. H.F.A was supported by Austrian Marshall Plan Scholarship. P.V. was supported by the ERC LipoCHEX (340896) and the FWF stand-alone grant LipoLUNG (P 30968).

## Author Contributions

H.F.A designed and performed the majority of the experiments. P.W.V performed intravenous injections of B16F10 cells. H.H. conducted bioinformatics analysis. J.F. & M.P. performed cell migration and sphere formation assays. G.H. and M.G.V.H provided substantial guidance and shared lab space and equipment for histology experiments. H.F.A and J.G.B-S constructed the study and H.F.A wrote the manuscript.

## Acknowledgments

We acknowledge the support of DK-MCD, NAWI Graz, constructive discussions with Elke Stadelmeyer, and the technical support of Thomas Schreiner, Wolfgang Krispel, and Silvia Schauer.

## Supplementary Figures

**Figure S1.**
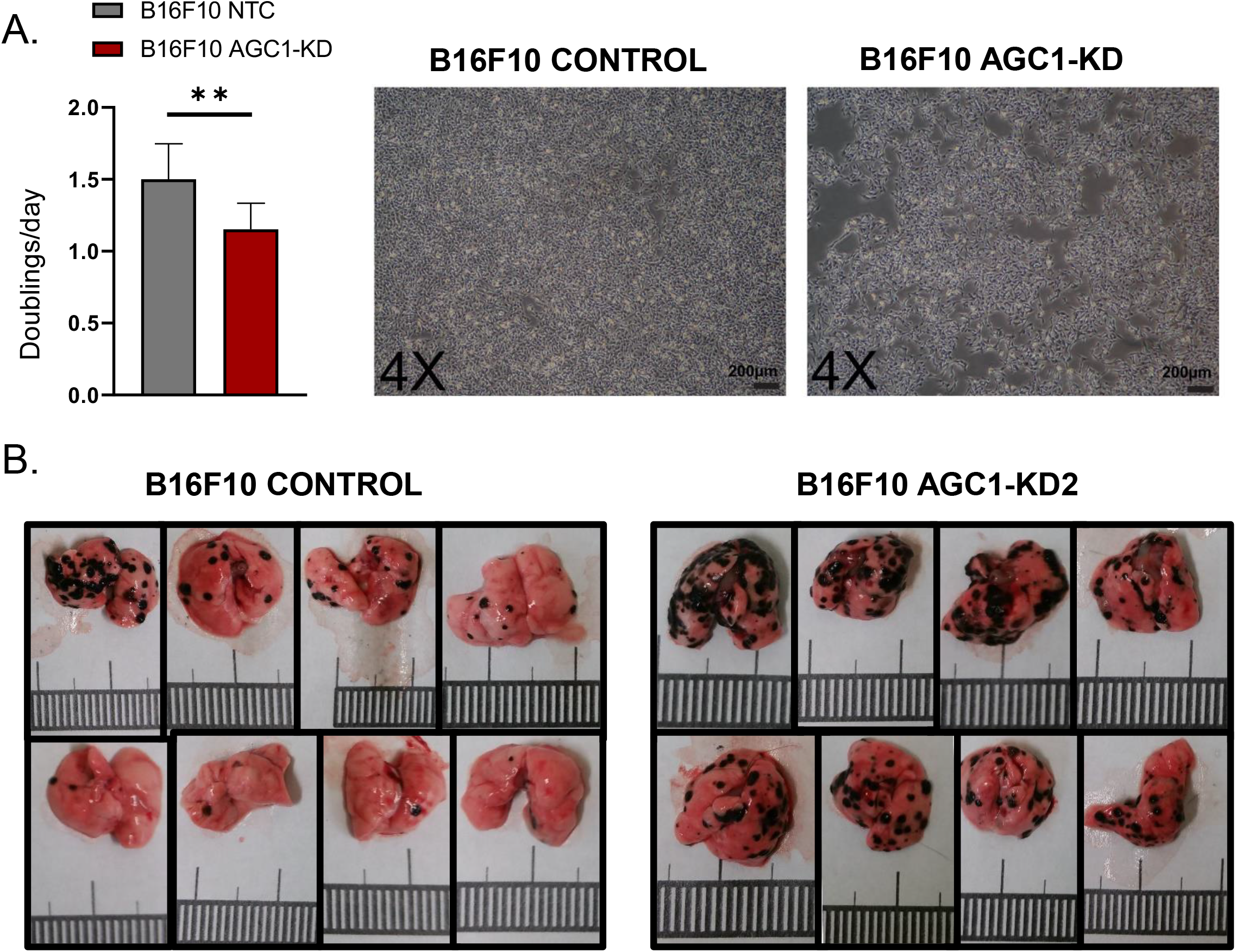
AGC1-knockdown in B16F10 impairs cell proliferation and increase lung metastasis. (A) (left) Proliferation rate of AGC1-KD and control (NTC) B16F10 cells in doublings/day, cultured in DMEM without pyruvate (n=3) means ±SDs are shown. (right) Representative pictures of B16F10 cells 64 hours after seeding for the proliferation assay. (B) Pictures of lungs with metastasis harvested from mice 16 days after control and AGC1-KD2 (sh908) B16F10 cells were injected intravenously. (related to Figure 2)

**Figure S2.**
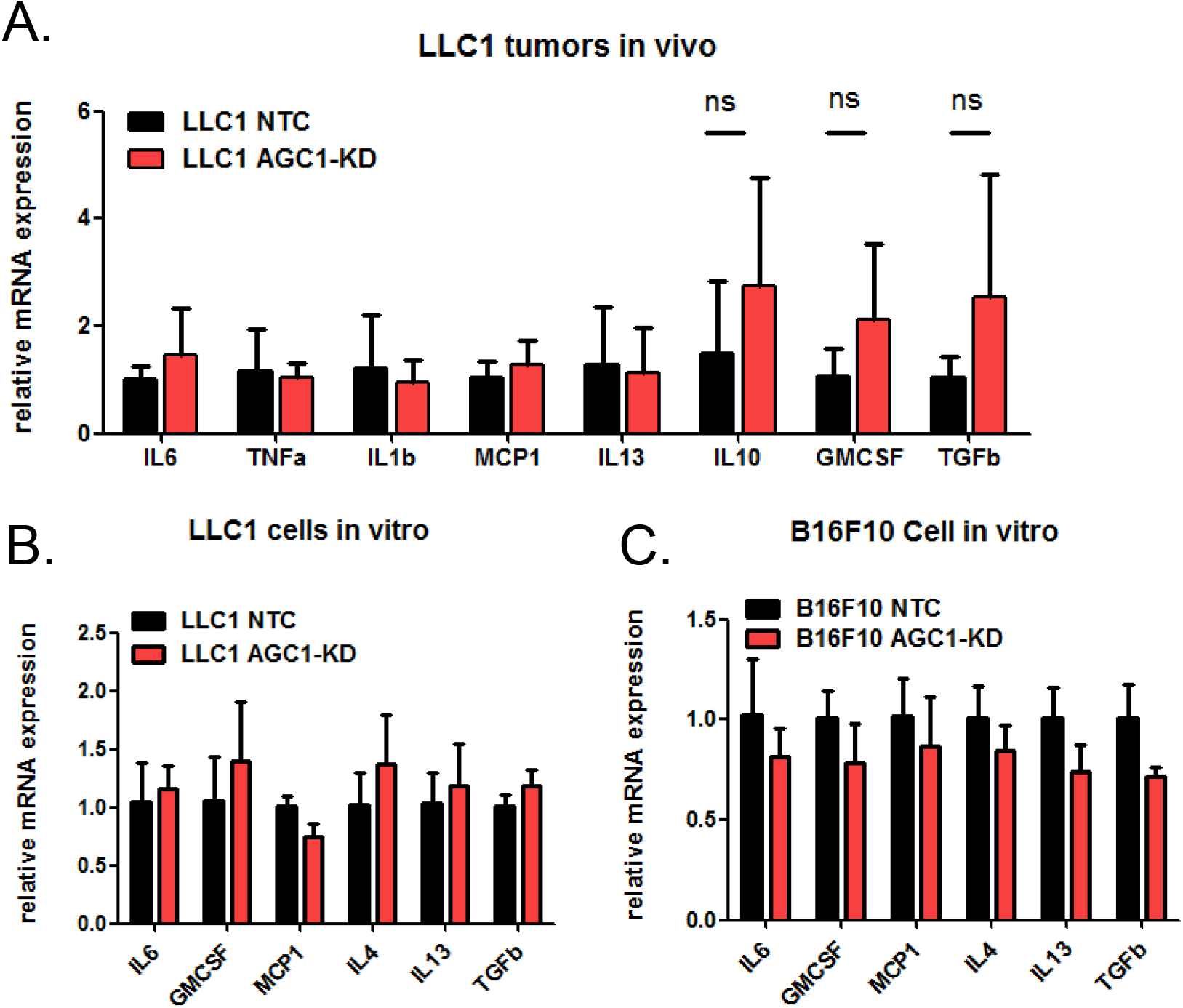
Levels of metabolites and inflammatory cytokines in AGC1-knockdown tumors. (A) Relative mRNA expressions of indicated genes in control (NTC) and AGC1-KD1 LLC1 tumors *in vivo* (n=6), means ±SDs are shown. (B-C) Relative mRNA expressions of indicated genes in control (NTC) and AGC1-KD1 (G) LLC1 and (H) B16F10 cells *in vitro* (n=3), means ±SDs are shown. Gene names; IL6: Interleukine-6, TNFa:Tumor necrosis factor alpha, IL1b:Interleukine1-beta, MCP1:Monocyte chemoattractant protein-1, IL4: Interleukine-4, IL13:Interleukine-13, IL10:Interleukine-10, GM-CSF:Granulocyte-macrophage colony-stimulating factor, and TGFb:Transforming growth factor beta. (related to Figure 4)

## Notes

### Competing Interest Statement

The authors have declared no competing interest.

